# Quantification of dengue virus specific T cell responses and correlation with viral load and clinical disease severity in acute dengue infection

**DOI:** 10.1101/325944

**Authors:** Dulharie T. Wijeratne, Samitha Fernando, Laksiri Gomes, Chandima Jeewandara, Anushka Ginneliya, Supun Samarasekara, Ananda Wijewickrama, Clare S. Hardman, Graham S. Ogg, Gathsaurie Neelika Malavige

## Abstract

**Background:** In order to understand the role of dengue virus (DENV) specific T cell responses that associate with protection, we studied their frequency and phenotype in relation to clinical disease severity and resolution of viraemia in a large cohort of patients with varying severity of acute dengue infection.

**Methodology/Principal findings:** Using ex vivo IFNγ ELISpot assays we determined the frequency of dengue viral peptide (DENV)-NS3, NS1 and NS5 responsive T cells in 74 adult patients with acute dengue infection and examined the association of responsive T cell frequency with the extent of viraemia and clinical disease severity. We found that total DENV-specific and DENV-NS3-specific T cell responses, were higher in patients with dengue fever (DF), when compared to those with dengue haemorrhagic fever (DHF). In addition, early appearance of DENV-specific T cell responses was significantly associated with milder clinical disease (p=0.02). DENV peptide specific T cell responses inversely correlated with the degree of viraemia, which was most significant for DENV-NS3 specific T cell responses (Spearman’s r = −0.47, p=0.0003). The frequency of T cell responses to NS1, NS5 and pooled DENV peptides, correlated with the degree of thrombocytopenia but had no association with levels of liver transaminases. In contrast, DENV-IgG inversely correlated with the degree of thrombocytopenia and levels of liver transaminases.

**Conclusions/significance:** Early appearance of DENV-specific T cell IFNγ responses appears to associate with milder clinical disease and resolution of viraemia, suggesting a protective role in acute dengue infection.

## Author summary

In order to understand the role of dengue virus (DENV) specific T cell responses in protection against infection, we studied T cell cytokine production in relation to clinical disease severity and resolution of viraemia in a large cohort of patients with varying severity of acute dengue infection. We found that DENV-specific T cell responses were higher in patients with dengue fever, when compared to those with dengue haemorrhagic fever. In addition, early appearance of DENV-specific T cell responses was significantly associated with milder clinical disease (p=0.02). DENV peptide specific T cell responses inversely correlated with the degree of viraemia, which was most significant for DENV-NS3 specific T cell responses (Spearman’s r = −0.47, p=0.0003). The frequency of NS1, NS5 and pooled DENV peptides, correlated with the degree of thrombocytopenia but had no association with liver transaminases. Our data suggest that early appearance of DENV-specific T cell IFNγ responses appear to associate with milder clinical disease and resolution of viraemia, suggesting a protective role in acute dengue infection.

## Introduction

Dengue virus is the cause of the most common mosquito-borne viral infection worldwide, indeed over half of the global population live in areas where there is intense dengue transmission putting them at risk of dengue infection [1]. Dengue virus causes 390 million infections annually, of which nearly a quarter are clinically apparent causing a spectrum of disease phenotypes ranging from mild dengue fever (DF) to dengue hemorrhagic fever (DHF). DHF is defined by a transient increase in vascular permeability resulting in plasma leakage, with high fever, bleeding, thrombocytopenia and haemoconcentration, which can lead to shock (dengue shock syndrome (DSS))[2]. It is however not fully understood why some people develop more severe forms of the disease, with patient history, immunity, age, viral serotype, sub-strain and epidemiological factors all postulated to play a role[3]. It was highlighted during a recent summit to identify correlates of protection for dengue, that dengue virus (DENV) specific T cell immunity should be studied in more detail, in order to develop safe and effective dengue vaccines[4].

Although a dengue vaccine (Denvaxia^®^) is now licensed in several countries, the efficacy is low in dengue seronegative individuals and provides only partial protection against DENV2 [5]. Although it is now generally believed that DENV specific T cells are protective, it is important that dengue vaccines should not induce “harmful” T cell immunity [4, 6–8]. Indeed, a significant hurdle in developing an efficacious dengue vaccine has been our limited understanding of the protective immune response in acute dengue infection and the added complexity of the presence of four DENV serotypes that are highly homologous. Seemingly conflicting evidence as to the role of antigen-specific T cells during dengue infection is reported in the literature.

T cell responses to DENV are predominantly directed towards the nonstructural proteins (NS), with the majority of the CD8+ T cell responses directed towards NS3 followed by NS5 and CD4+ T cell responses to envelope, PrM and NS1 proteins [9–11]. It was believed that highly cross-reactive T cells specific to DENV-NS3, and other proteins, associate with severe clinical disease (DHF), and it was thought that these cells contribute to DHF by inducing a ‘cytokine storm’[12–15]. It is hypothesized in the ‘original antigenic sin’ theory that T cell responses against the initial DENV serotype of primary infection persist and dominate during subsequent infections; and that these T cells are suboptimal in inducing robust antiviral responses upon re-challenge [13, 14, 16]. However, it has been shown that DENV-NS3 specific T cell responses were at very low frequency during acute disease, and only detected in the convalescent phase pointing away from a role in vascular leak [14, 16, 17]. Recently it was observed that DENV-specific T cells are found in large numbers in the skin during acute dengue infection, and it is speculated that highly cross-reactive, pathogenic skin T cells could be contributing to DHF, despite being absent or present at low frequencies in the peripheral blood [8, 18]. As the frequency of skin resident DENV-specific T cells was investigated in a small patient cohort, it is not yet clear whether the frequency of the skin T cells associated with clinical disease severity.

Conversely some studies in both humans and mouse models have shown that DENV-specific T cells in the blood are likely to be protective [19–23]. It was shown that in individuals who were naturally infected with DENV, polyfunctional CD8+ T cells responses of higher magnitude and breadth were seen for HLA alleles associated with protection [21]. Similar findings were seen with DENV-specific CD4+ T cell responses [23]. Our previous studies have also shown that the magnitude of IFNγ-producing DENV NS3-specific memory T cell responses was similar in those who had varying severity of recovered past dengue infection, suggesting that the magnitude of the memory T cell response does not correlate with clinical disease severity[22]. While many studies have been carried out to elucidate the functionality of T cell responses in dengue, these have been limited to studying T cells specific for particular HLA types by using tetramers/pentamers [16, 18], or to investigating T cell responses in individuals with unknown severity of dengue.

To aid the generation of an effective vaccine it will be important to understand the role, phenotype and frequency of dengue-specific T cell responses in relation to clinical disease severity and clearance of viraemia [6, 7]. Therefore, here we investigate T cell responses to immunodominant DENV NS proteins in patients with DHF and DF, and analyse the association of such responses with resolution of viraemia.

## Methods

### Recruitment of patients for analysis of the functionality of T cell responses

We recruited 74 adult patients with acute dengue infection from the National Infectious Diseases Institute, between day 4 - 8 of illness, following informed written consent. All clinical features were recorded several times each day, from time of admission to discharge. Ultra sound scans were performed to determine the presence of fluid leakage in pleural and peritoneal cavities. Full blood counts, and liver transaminase measurements were performed serially through the illness. Clinical disease severity was classified according to the 2011 WHO dengue diagnostic criteria [24]. Accordingly, patients with ultrasound scan evidence of plasma leakage (those who had pleural effusions or ascites) were classified as having DHF. Shock was defined as having cold clammy skin, along with a narrowing of pulse pressure of ≤ 20 mmHg. Based on this classification, 45 patients had DHF and 29 patients had dengue fever (DF) of the 74 patients recruited for the study.

### Ethics statement

The study was approved by the Ethical Review Committee of The University of Sri Jayewardenepura. All patients were adults and recruited post written consent.

### Serology

Acute dengue infection was confirmed in serum samples using a PCR (see below) and dengue antibody detection. Dengue antibody assays were completed using a commercial capture-IgM and IgG ELISA (Panbio, Brisbane, Australia) [25, 26]. Based on the WHO criteria, those who had an IgM: IgG ratio of >1.2 were considered to have a primary dengue infection, while patients with IgM: IgG ratios <1.2 were categorized under secondary dengue infection [27].

The DENV-specific IgM and IgG ELISA was also used to semi-quantitatively determine the DENV-specific IgM and IgG titres, which were expressed in PanBio units.

### Qualitative and quantitative assessment of viral loads

DENV were serotyped and viral titres quantified as previously described [28]. RNA was extracted from the serum samples using QIAamp Viral RNA Mini Kit (Qiagen, USA) according to the manufacturer’s protocol. Multiplex quantitative real-time PCR was performed as previously described using the CDC real time PCR assay for detection of the dengue virus [29], and modified to quantify the DENV. Oligonucleotide primers and a dual labeled probe for DENV 1,2,3,4 serotypes were used (Life technologies, India) based on published sequences [29]. In order to quantify viruses, standard curves of DENV serotypes were generated as previously described in Fernando, S. *et.al* [28].

### Peptides

The peptide arrays spanning DENV NS1 (DENV-2 Singapore/S275/1990, NS1 protein NR-2751), NS3 (DENV-3, Philippines/H87/1956, NS3 protein, NR-2754) and NS5 proteins (DENV-2, New Guinea C (NGC), NS5 protein, NR-2746) were obtained from the NIH Biodefense and Emerging Infections Research Resource Repository, NIAID, NIH. The DENV NS3 peptide array consisted of 105, 4-17 mers peptides, NS1 and NS5 proteins were comprised of 60 and 156 peptides respectively. The peptides were reconstituted as previously described [30]. NS1, NS3 and NS5 peptides were pooled separately to represent the DENV-NS1, NS3 and NS5 proteins. In addition, total NS1, NS3 and NS5 peptides were combined to represent a ‘DENV-all’ pool of peptides.

### *Ex vivo* ELISpot assay

Ex vivo IFNγ ELISpot assays were carried out as previously discussed using freshly isolated peripheral blood mononuclear cells (PBMC) obtained from 74 patients [22]. DENV-NS3, NS1, NS5 and the combined DENV-ALL peptides were added at a final concentration of 10 μM and incubated overnight as previously described [16, 31]. All peptides were tested in duplicate. PHA was included as a positive control of cytokine stimulation and media alone was applied to the PBMCs as a negative control. The spots were enumerated using an automated ELISpot reader (AID Germany). Background (PBMCs plus media alone) was subtracted and data expressed as number of spot-forming units (SFU) per 10^6^ PBMCs.

### Quantitative cytokine assays

Quantitative ELISA for TNFα (Biolegend USA) and IL-2 (Mabtech, Sweden) were performed on ELISpot culture supernatants according to the manufacturer’s instructions.

### Statistical analysis

PRISM version 6 was used for statistical analysis. As the data were not normally distributed, differences in means were compared using the Mann-Whitney U test (two tailed). Spearman rank order correlation coefficient was used to evaluate the correlation between variables.

## Results

### Patient clinical and laboratory features

To investigate the role of T cells in the progression of dengue infection we stratified patients based on disease severity. The clinical and laboratory features of the 74 patients recruited to the study are shown in table 1. Of these 74 patients, 45 had DHF and 29 had DF, and all 45 patients with DHF had ascites with 10 of them also experiencing pleural effusions. None of the patients developed shock and only one person progressed to bleeding manifestations (table 1). The median duration of illness when recruited to the study was similar for patients with DF (median 5, IQR 5 to 6 days) and DHF (median 5, IQR 4 to 6 days).

**Table 1:**
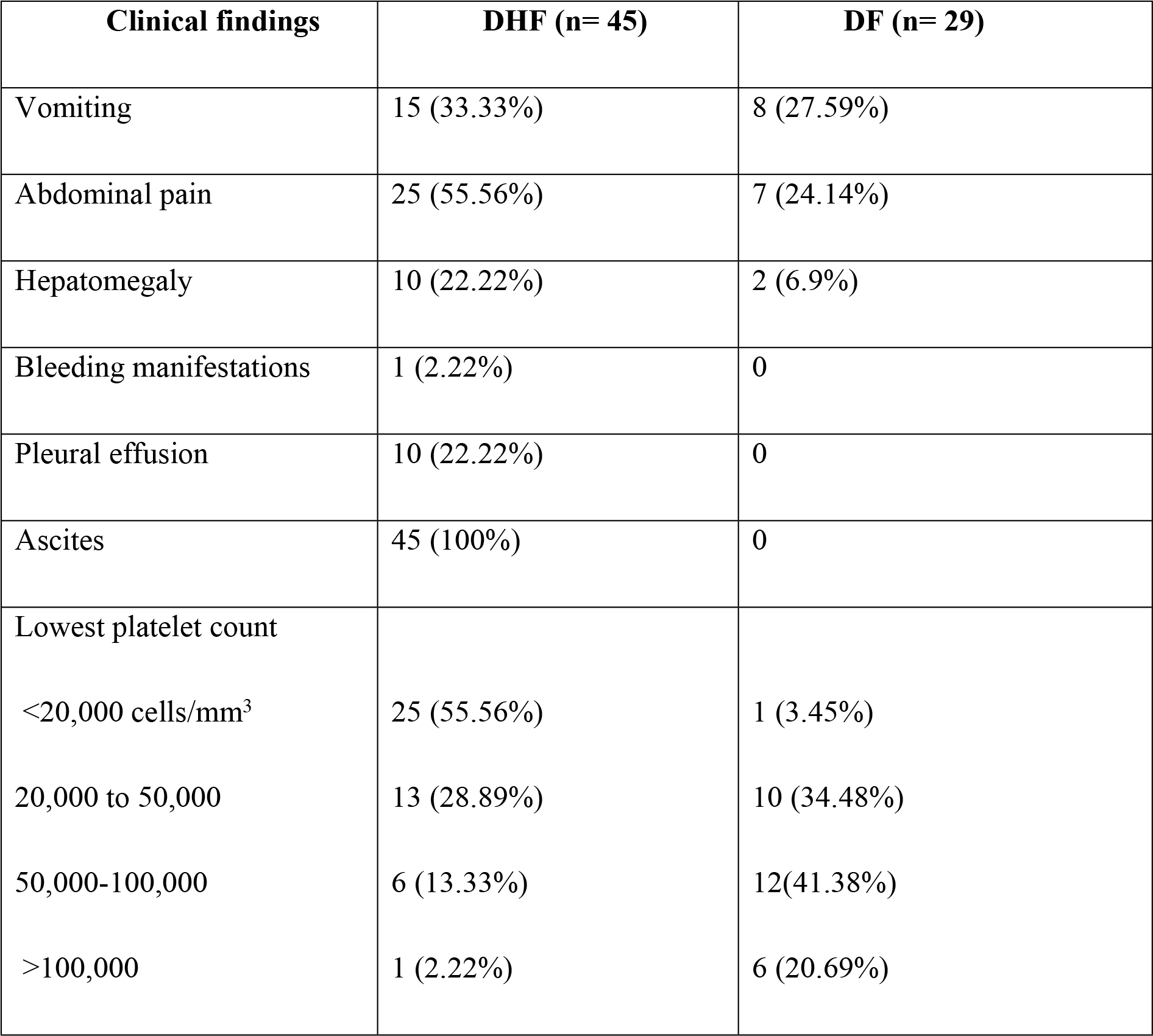
Clinical and laboratory characteristics of patients with DHF and DF.

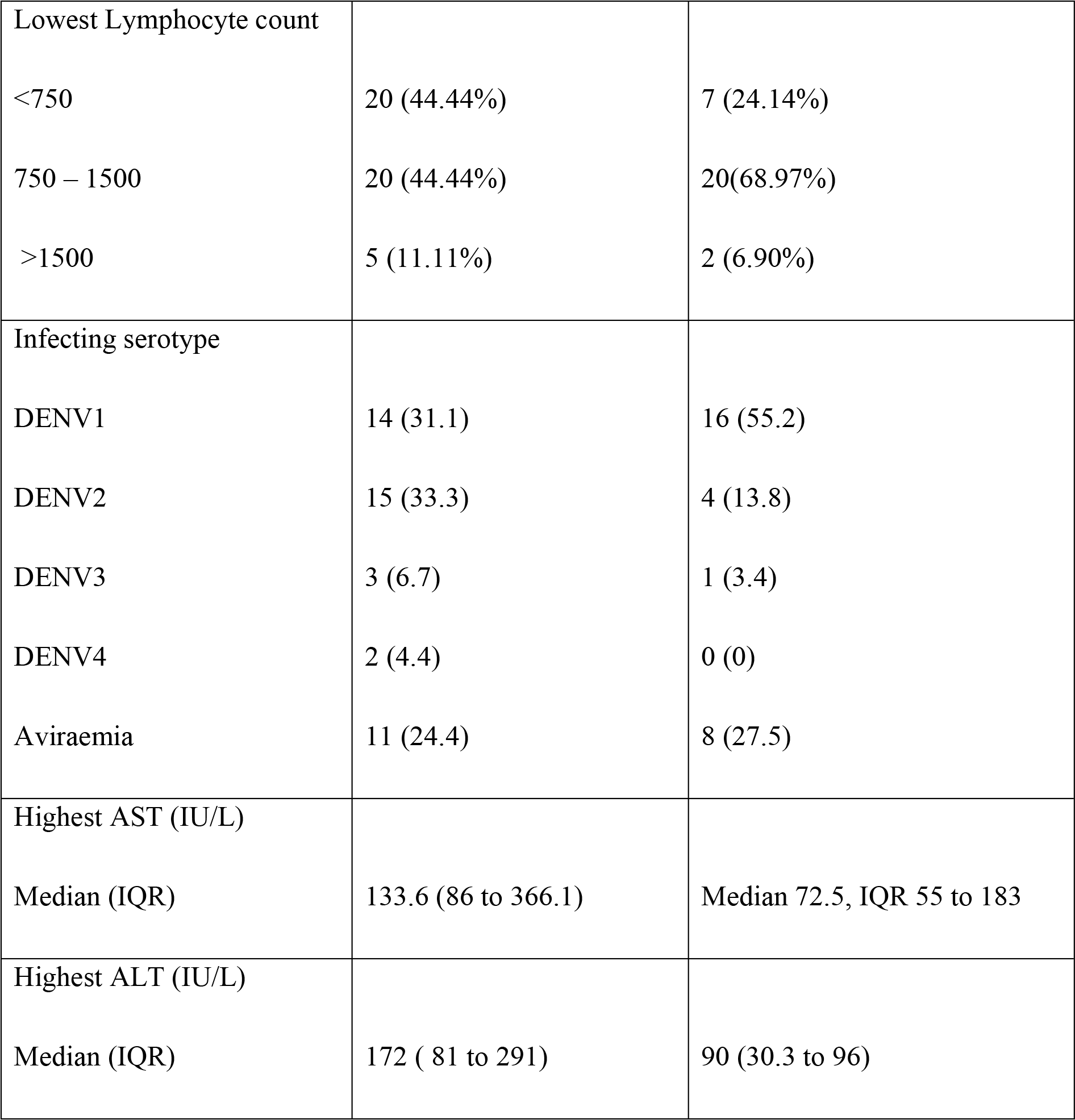

### Ex vivo IFNγ responses in patients with acute dengue infection

To evaluate the role of T cell derived cytokine in the immunopathology or regulation of acute dengue infection, we stimulated PBMCs isolated from patients with either DF or DHF with peptides constituting DENV-derived non-structural protein (NS) and assessed cytokine production by ELISPOT. We stimulated the patient PBMCs with different pools of overlapping peptides making up either full length NS1, NS3 or NS5 protein, or a pool of total NS1, 3 and 5 peptides (DENV-all). NS3 and NS5 were selected for investigation as CD8+ T cell responses have been shown to be directed to these proteins and CD4+ T cells have been shown to target structural proteins and NS1 as the main non-structural protein[11, 21]. This combination of NS proteins from the particular DENV strains has previously been used to study DENV specific T cell responses [32]. We used this ex vivo ELISPOT method to model antigen presentation of dengue-derived peptides to antigen-specific T cells in vitro and assessed T cell activation by IFN<u>γ</u> production, as a representative cytokine produced by T cells during dengue infection. T cell responses to the pooled DENV peptides (DENV-ALL) (p=0.02) were higher in PBMCs derived from patients with DF than DHF patients and the NS3-specific responses showed a trend to be higher in those with DF than DHF (Fig. 1A). T cell responses to DENV-NS1 peptides were similar in patients with DF and DHF. We did not detect TNFα in the ex vivo ELISpot culture supernatants, which is in contrast to studies performed by others on T cell clones that implied TNFα producing DENV-specific T cells contribute to disease pathogenesis [15]. We also did not detect significant quantities of IL-2.

**Figure 1:**
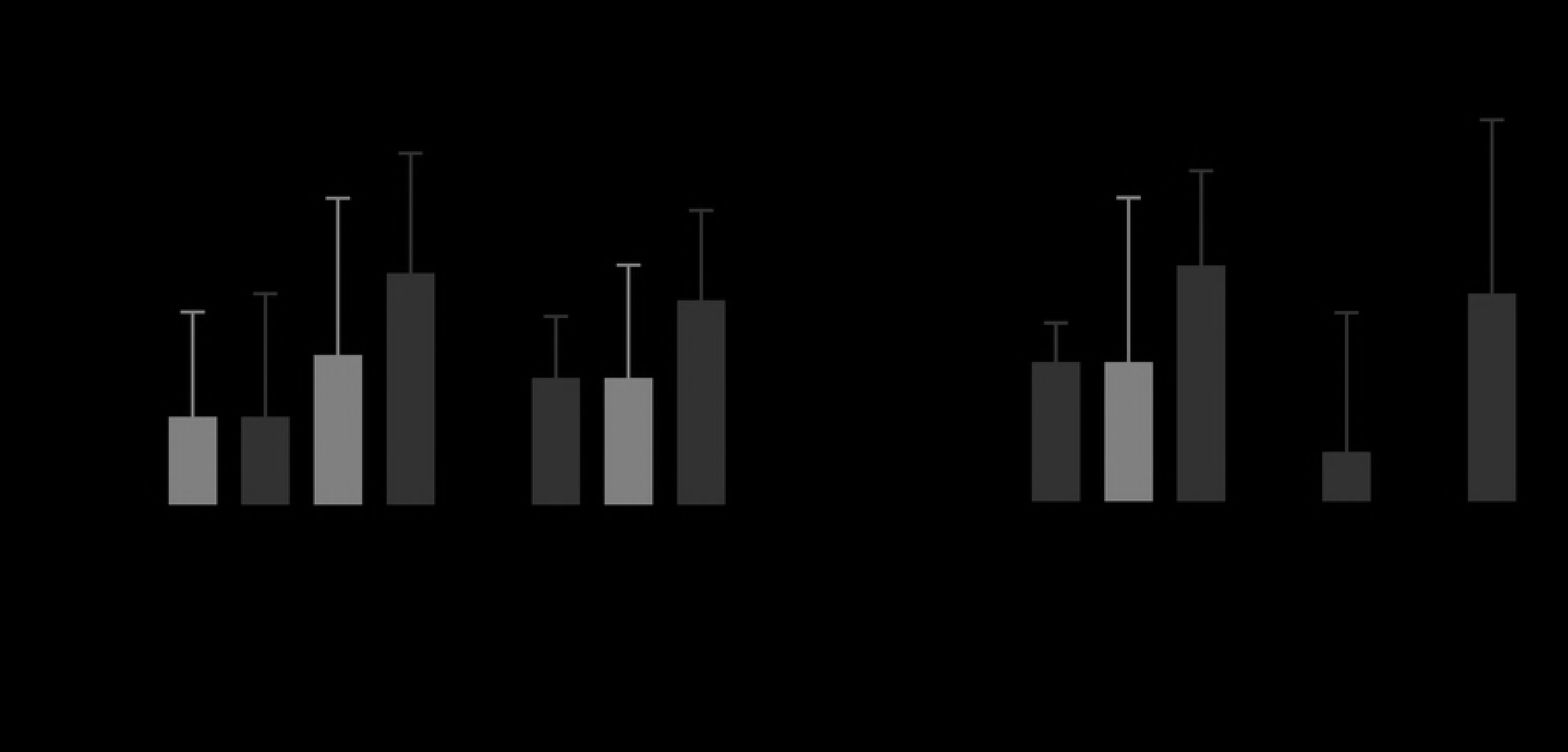
Ex vivo ELISpot responses to DENV peptides in patients with DHF and DF. A) Ex vivo IFN<u>γ</u> ELISpot responses to DENV NS1, NS3, NS5 and combined DENV overlapping peptides in patients with DF (n=29) and DHF (n=45). (B) Ex vivo IFN<u>γ</u> ELISpot responses to DENV peptides in patients who were recruited on day 4 since onset of illness with DF (n=6) and DHF (n=12). Error bars represent the median and the interquartile range. *P<0.05.

To assess if detection of DENV specific T cell responses before the onset of the critical phase (vascular leakage phase), was associated with a reduced likelihood of developing leakage, we isolated the data from DF and DHF patients recruited on day 4 post the onset of illness and analysed IFNγ production by peptide stimulated PBMCs. None of the patients had evidence of vascular leakage on day 4 of illness and those who developed leakage (patients with DHF), did so on day 5 or 6. DF patients had a significantly higher IFNγ secretion response (p=0.02) to the DENV-all peptide pool (median 42.5, IQR=22.5 to 945 SFU/10^6^ PBMCs), when compared to DHF patients (median 0, IQR=0 to 12.5 SFU/10^6^ PBMCs) (Fig 1B). As such, significantly higher DENV-specific T cell responses were seen in those who did not develop fluid leakage, and those who had lower DENV-specific T cell responses proceeded to develop fluid leakage (DHF). Responses to DENV-NS3, NS1 and NS5 also appeared higher in patients with DF at this time point, although this did not reach statistical significance (Fig. 1B).

To further assess the time-course of the response we obtained a second blood sample from eight patients within our cohort two days after collection of the first sample. T cell responses to DENV-ALL and DENV-NS3 peptides increased from the first sample (day 4) to the second (day 6), but it was not statistically significant (p>0.05) (Supplementary fig. 1).

### Laboratory parameters and DENV-specific T cell responses

Thrombocytopenia is associated with clinical disease severity and higher degrees of thrombocytopenia are seen in those with DHF compared to those with DF[24]. We found that DENV peptide specific T cell responses inversely correlated with the degree of thrombocytopenia. While this inverse correlation with T cell responses and platelet counts was significant for DENV-NS1 (Spearmans r=0.26, p=0.01) (Fig 2A), NS5 (Spearmans r=0.4, p=0.0002) (Fig 2B) and DENV-All (Spearmans r=0.31, p=0.005) (Fig 2C), it was not significant for NS3 (Spearmans r=0.18, p=0.09) (Fig 2D). No association was seen with DENV-peptide specific T cell responses and aspartate transaminase (AST) and alanine transaminase (ALT) (Supplementary fig 2), which are indicators of liver dysfunction [28, 33].

**Figure 2:**
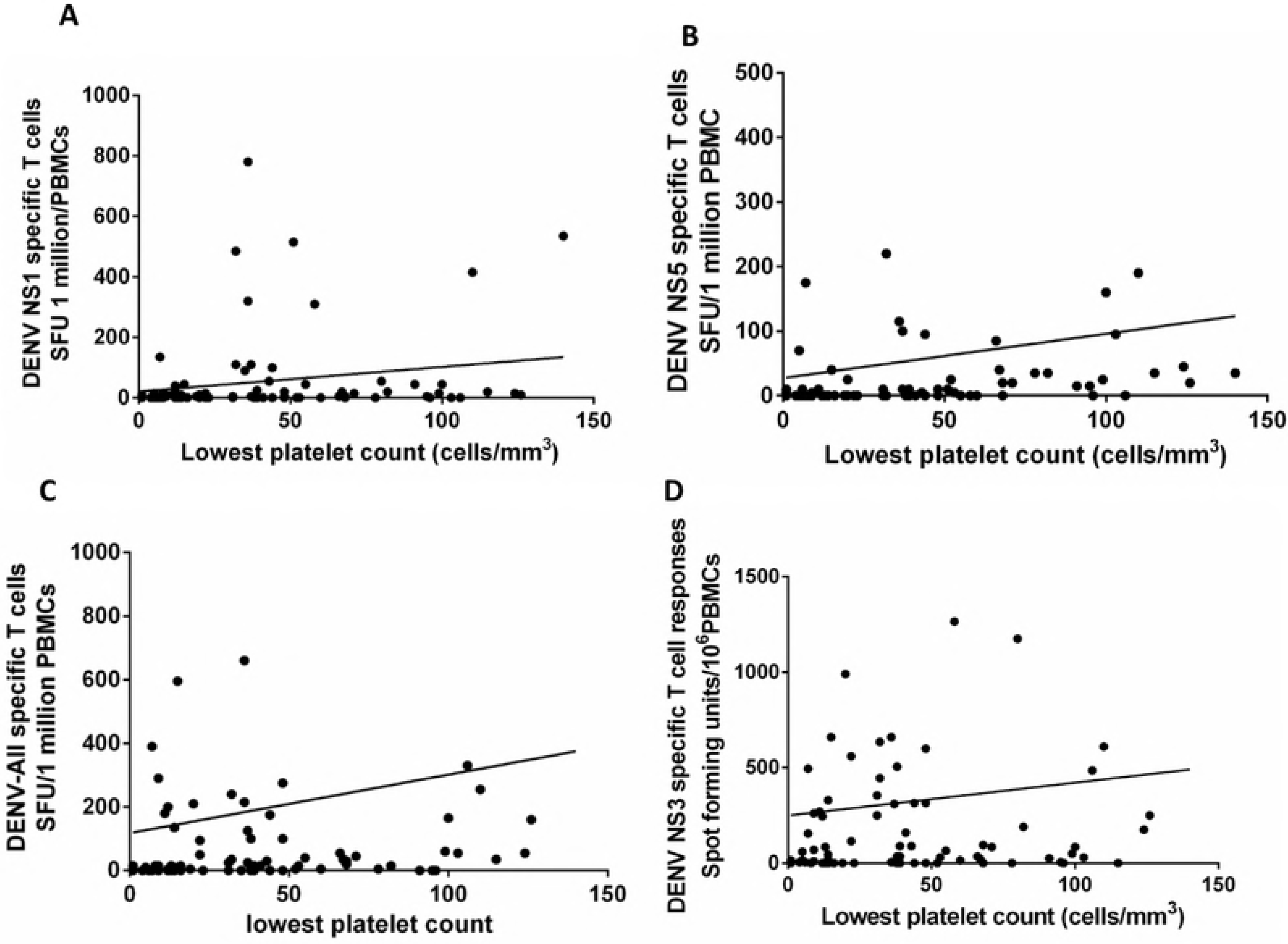
Relationship between laboratory parameters and DENV specific T cells in patients with acute dengue. Platelet counts were correlated with ex vivo IFNγ ELISpot responses to DENV NS1 (Spearmans r=0.26, p=0.01) (A), NS5 (Spearmans r=0.4, p=0.0002) (B) and the overall DENV (Spearmans r=0.31, p=0.005) (C) and NS3 (Spearmans r=0.18, p=0.09) (D) overlapping peptides in patients (n=74).

### The relationship between DENV serotype and T cell responses

While some studies report that certain DENV serotypes associate with DHF [34, 35], others have shown that the risk of DHF is similar regardless of serotype [36]. Therefore, we proceeded to determine whether there were differences in the T cell responses to DENV-proteins based on the viral serotype that the patients were infected with. Within our cohort 30 (40.5%) patients were infected with DENV1, 19 (25.7%) with DENV2, 4 (5.4%) with DENV-3 and 2 (2.7%) with DENV-4 (Table 1). The serotype could not be determined in 19 (25.7%) patients, as they were not viraemic at the time of recruitment. DHF developed in 14/30 (46.7%) of the patients infected with DENV-1 and 15/19 (78.9%) of those infected with the DENV-2 and in 11/19 (57.9%) who were aviraemic at the time of recruitment (Fig 3A). Thus, it appeared as if DENV-2 infection was more likely to lead to development of DHF (odds ratio 3.3, 95% CI 0.93 to 12.1), however the association was not statistically significant (p=0.08) in this cohort. Aviraemic individuals displayed significantly higher IFNγ T cell responses to NS1 (p=0.002), NS3 (p=0.02), NS5 (p=0.02) and DENV-ALL pooled peptides (p=0.0004) when compared to those who were viraemic at the time of recruitment (Fig 3B). In addition, those who were infected with the DENV-2 serotype, with a trend towards increased DHF susceptibility, had significantly lower responses to NS1 (p=0.002), NS3 (p=0.04), NS5 (p=0.003) and DENV-All (p=0.0003) peptides when compared to those who were infected with DENV-1.

Multiple alignment of the NS5 protein sequences of DENV2 (58 sequences) and DENV3 (28 sequences) was performed using virus variation resource [37] and analysed using Clustal omega and showed a sequence identity of > 72.1% between the NS5 proteins of these viral serotypes [38]. Multiple alignment of the NS3 protein of DENV2 (61 sequences) and DENV3 (28 sequences) showed a sequence identity of > 72.02%. Multiple alignment of the NS1 protein of DENV2 (62 sequences) and DENV3 (28 sequences) showed a sequence identity of >65.11%. The homology between DENV1 and DEN2 NS5 was >71.83% (comparison of 102 DENV1 sequences and 58 DENV2 sequences) [37], while the homology between DENV1 and DENV3 NS3 was >76.9% (comparison of 102 DENV1 sequences and 28 DENV3 sequences) [37]. Therefore, the differential response to DENV serotype is unlikely to be profoundly influenced by a difference in NS1 and NS2 protein sequences.

**Figure 3:**
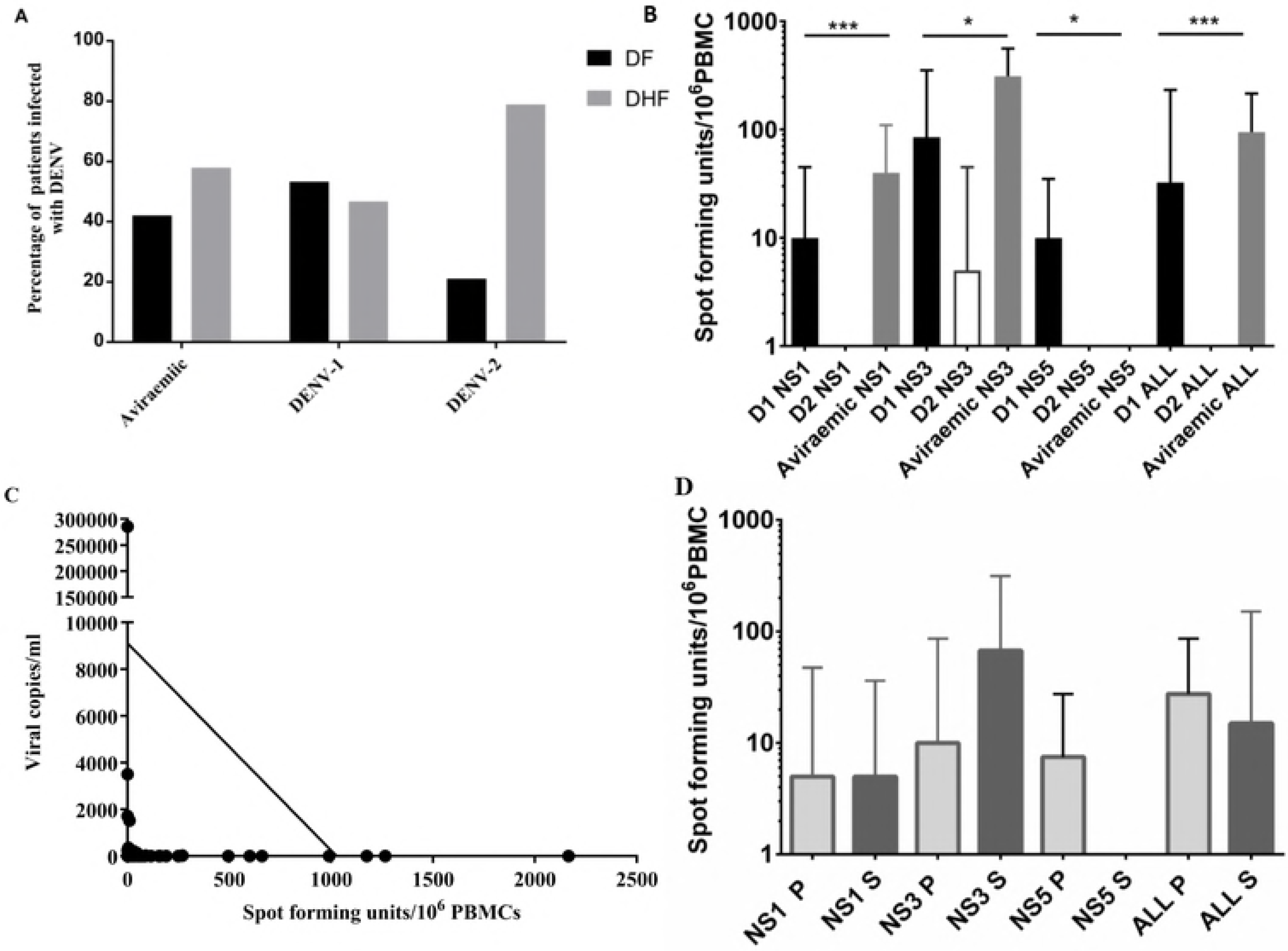
Relationship between DENV serotype, clinical disease severity, viraemia and T cell responses. A: Proportion of patients, infected with DENV1 (n=30), DENV2 (n=19) or were aviraemic (n=19) who developed DF or DHF. B: Ex vivo ELISpot responses to DENV NS1, NS3, NS5 and combined DENV-ALL overlapping peptides in patients who were infected with DENV-1 (n=30), DENV-2 (n=19) or who were aviraemic (n=19). Error bars represent the median and the interquartile range. **P<0.05, **P<0.01, ***P<0.001 (C) Correlation between DENV NS3-specific T cell responses and degree of viraemia (Spearman’s r = −0.47, p=0.0003). (D) Ex vivo ELISpot responses to DENV NS1, NS3, NS5 and the overall DENV overlapping peptides in patients who had primary dengue (P) (n=19) and secondary dengue infection (S) (n=48)

### Viraemia and DENV specific T cell responses

DHF patients have been shown to have higher viral loads, exhibit prolonged viraemia [39, 40] and persistent DENV-NS1 antigenaemia [41, 42]. As such we attempted to elucidate a correlation between T cell cytokine responses and viremia. DENV specific T cell responses to NS1, NS3 and NS5 peptides in addition to the pooled peptides (DENV-ALL) inversely correlated with the degree of viraemia, which was most significant for DENV-NS3 specific T cell responses (Spearman’s r = -−0.47, p=0.0003) (Fig. 3C and supplementary Fig 3). The viral loads significantly inversely correlated with the platelet counts (Spearmans r=−0.34, p=0.01), with the platelet counts being lowest in individuals with the highest viral loads (data not shown).

It is thought that a second dengue virus infection with a different viral serotype is a risk factor for developing DHF[43]. To determine the effect of secondary infection on the resulting T cell response, we characterized patient infection history and assayed patient blood for the presence of dengue specific IgM and IgG. Primary infection was defined by DENV-specific IgM:IgG >1.2 [24]. Accordingly, 19 (25.7 %) patients were classified as experiencing a primary dengue infection and 48 (64.9%) were defined as secondary dengue infection. The antibody results were inconclusive for 7 (9.4%) patients. Our results showed no significant difference in DENV specific T cell responses between primary and secondary dengue infection patient groups (p>0.05) for any of the DENV peptide pools (Fig. 3D).

We semi-quantitatively determined the DENV-specific IgM and IgG antibody titres in all patients with DF and DHF, and we found that neither the DENV-IgM nor IgG antibody titres correlated with T cell responses to DENV-NS1, NS5 and NS3. However, the DENV-specific IgG antibody titres inversely correlated with viral loads in those with DHF (Spearman’s r=-0.37, p=0.03) (Fig. 4A), but not in those with DF (Spearmans r=-0.25, p=0.16) (data not shown). In analysis of the IgG antibody titres of all patients (n=74) they too inversely correlated with the degree of thrombocytopenia (Spearmans r=-0.29, p=0.009) (Fig. 4B). DENV-specific IgG also correlated with the highest aspartate transaminase (AST) (Spearmans r=0.51, p=0.004) (Fig. 4C) and alanine transaminase (ALT) levels (Spearmans r=0.4, p=0.03) in all patients with acute dengue infection (Fig. 4D).

**Figure 4:**
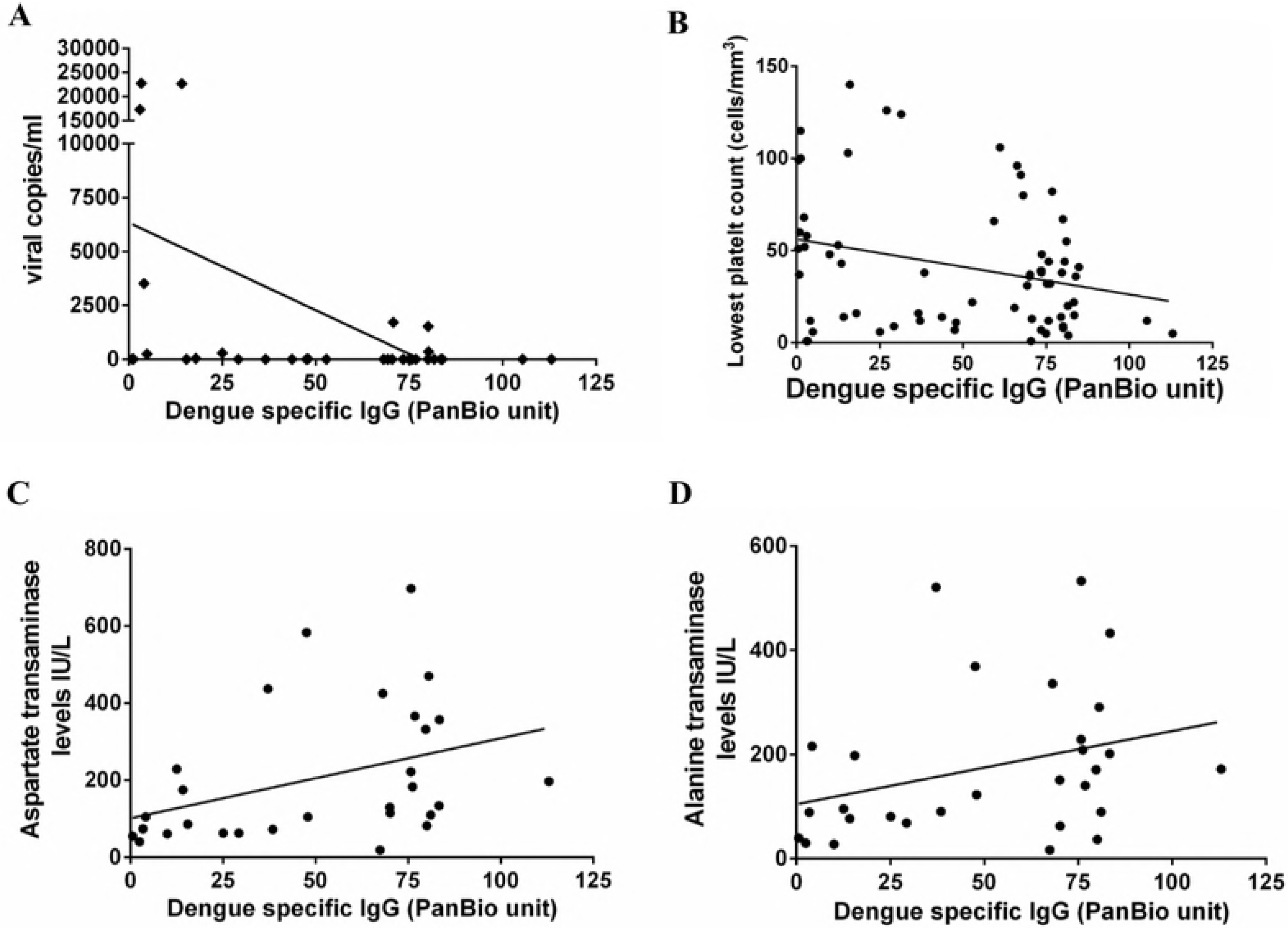
DENV-specific IgG responses and laboratory parameters of clinical disease severity in patients with acute dengue. Analysis of the correlation between DENV-specific IgG responses and (A) degree of viraemia (Spearmans r=-0.25, p=0.16) in patients with DHF, (B) degree of thrombocytopenia (Spearmans r=-0.29, p=0.009), (C) aspartate transaminase (AST) levels (Spearmans r=0.51, p=0.004) and (D) alanine transaminase (ALT) levels (Spearmans r=0.51, p=0.004) in all patients (n=74) with acute dengue infection.

## Discussion

In this study we set out to investigate the role of T cells in dengue immunity and found that DENV-specific T cells are present at low frequency during acute infection, consistent with previous reports published by us and by others [16, 17, 44]. IFNγ production was significantly higher in patients with DF as opposed to DHF, especially during early infection. Patients who had higher DENV-NS3 specific T cell responses on day four since the onset of illness (before development of fluid leakage), were significantly more likely to develop DF than DHF. In addition, the frequency of pooled DENV-peptide specific, in particular DENV-NS3 specific, T cell responses was associated with resolution of viraemia. Aviraemic patients had significantly higher DENV-specific T cell responses when compared to those who were viraemic. T cell IFNγ responses to DENV NS1, NS5 and pooled DENV (NS1, NS3 and NS5) peptides inversely correlated with the degree of thrombocytopenia, but we did not show any relationship with liver transaminases (AST and ALT levels). Both the degree of thrombocytopenia and a rise in both AST and ALT, have been shown to associate with dengue severity [24, 28]. Therefore, our data show that the early appearance of DENV-NS3 specific T cell responses is associated with milder disease, which is compatible with recent studies regarding the role of T cells in DENV infection [18, 21–23]. This suggests that DENV-peptide specific T cells are protective against developing severe forms of dengue infection.

Although Appana et al also evaluated *ex vivo* IFNγ to selected peptides of structural and non-structural DENV proteins by ELISpot assays, they did not find any differences in the frequency of DENV-specific T cell responses in patients with DF when compared to those with DHF [45]. However, only peptides that were predicted to bind to certain major HLA alleles were included in the authors’ peptide pools used in the ELISpot assays [45], whereas here we utilised peptides spanning the entire length of DENV NS1, NS3 and NS5, proteins. This difference in experimental approach may have affected the cytokine production profile of the responding T cells in the different disease states. In addition, as the viability and function of T cells have been shown to be affected in those with acute dengue infection [46], we used freshly isolated PBMCs in all our experiments to limit extraneous cellular stress in contrast to previous studies [15, 21, 45, 47].

In general, more severe forms of dengue infection are observed during a secondary heterologous dengue infection [4], which gave rise to the hypothesis that cross reactive T cells responding to the primary infecting DENV serotype are suboptimal in clearing the secondary virus, and lead to development of more severe disease [14, 16]. In these studies, it was shown that a tetramer of different viral specificity to the current infecting DENV serotype, sometimes had a higher affinity to the DENV specific T cells [16]. In our study, we did not observe any difference in IFNγ production in overall *ex vivo* ELISpot assays from PBMCs derived from patients with primary or secondary dengue infection; however, we did not examine variant peptide-specific responses. The broad differences we observed in DENV-specific T cell responses correlated only with clinical disease severity. Interestingly, the DENV-specific IgG levels, which were measured semi-quantitatively, inversely correlated with the degree of thrombocytopenia and also AST and ALT levels, which are known to associate with liver damage. DENV-specific IgG levels are known to be significantly higher in patients with secondary dengue, compared to primary dengue, indeed it is one of the criteria for definition of a secondary dengue infection. Therefore, our data show that antibodies may contribute to severe disease, particularly during secondary dengue infection.

In summary, we found that DENV-specific T cell IFNγ responses, were associated with milder clinical disease severity and resolution of viraemia, suggesting a protective role for peptide specific T cells early in acute dengue infection.

## Supporting information captions

**Supplementary figure 1**: Frequency of DENV specific NS1, NS3, NS5 and All overlapping peptide responses in patients with acute dengue (n=8) on day 4 and day 6 of illness.

**Supplementary figure 2**: Association of DENV specific NS1, NS3, NS5 and All overlapping peptide responses in patients with acute dengue (n=74) with the highest recorded aspartate transaminase level (NS1 Spearnman’s r=0.15, p=0.4; NS3 Spearman’s r=0.34, p=0.05; NS5 Spearman’s r=0.16, p=0.37; All Spearman’s r=0.33, p=0.06) (A) and alanine transaminase level (NS1 Spearman’s r=12, p=0.5; NS3 Spearman’s r=0.25, p=0.15; NS5 Spearman’s r=0.19, p=0.29; All Spearman’s r=0.31, p=0.08) (B)

**Supplementary figure 3: Relationship between viraemia and DENV T cell responses**

(A) Correlation between DENV All-Specific T cell responses and degree of viraemia (Spearman’s r = −0.38, p=0.004).

(B) Correlation between DENV NS5-Specific T cell responses and degree of viraemia (Spearman’s r = −0.28, p=0.04).

(C) Correlation between DENV NS1-Specific T cell responses and degree of viraemia (Spearman’s r = −0.31, p=0.02).

